# PDL1 CHECKPOINT BLOCKADE SYNERGIZES WITH NILOTINIB BUT NOT DASATINIB TO PREVENT LEUKEMIA RELAPSE

**DOI:** 10.1101/2025.07.02.662848

**Authors:** Elizabeth C. Morgan, Hrishi Venkatesh, Enoc Granados Centeno, Lynn M. Heltemes-Harris, Gregory Hubbard, Taylor L. Maiers, Thamotharampillai Dileepan, Sean I. Tracy, Michael A. Farrar

## Abstract

Dasatinib and nilotinib are front line Tyrosine Kinase Inhibitors (TKIs) used to treat BCR-ABL+ B cell acute lymphoblastic leukemia (B-ALL) and BCR-ABL+ chronic myelogenous leukemia. We previously showed that combining nilotinib with anti-PD-L1 blockade significantly reduced leukemia relapse. The TKI dasatinib is more commonly used to treat to B-ALL. However, unlike nilotinib, dasatinib also inhibits SRC-family kinases, which may make it less efficacious in combination with anti-PDL1 blockade. Herein we assess the impact of nilotinib versus dasatinib on anti-leukemia immune responses. Dasatinib, but not nilotinib, inhibited T cell proliferation at high doses *in-vitro*, but neither TKI significantly impacted T cell function or expansion in response to immunization *in-vivo* with a model antigen (2W1S) plus polyIC-adjuvant. Dasatinib and nilotinib both reduced leukemic blasts equivalently after 5 days of treatment. In contrast, nilotinib and PD-L1 blockade, but not dasatinib plus anti-PDL1, prevented relapse several weeks later. Thus, dasatinib negatively impacts protective anti-leukemia T cell responses that prevent leukemia relapse.

## INTRODUCTION

BCR-ABL+ B-ALL leukemia is characterized by fusion of chromosomes 9 and 22 that results in the generation of the BCR-ABL oncogene, a constitutively active tyrosine kinase that can be inhibited by a variety of <u>T</u>yrosine <u>K</u>inase Inhibitors (TKIs) including imatinib, dasatinib, and nilotinib^1^. TKIs are extremely effective at treating patients with BCR-ABL+ CML. In contrast, while these TKIs are effective at inducing short-term remission in BCR-ABL+ B-ALL patients, these patients typically relapse with well characterized escape mutations in their tyrosine kinase domain^2^. B cell leukemias have relatively few non-synonymous mutations compared to other cancers such as melanoma or lung cancer. Nevertheless, robust anti-leukemia T cells responses can be observed in patients with B-ALL^3^. These findings suggest that anti-leukemia immune responses can be effective at targeting B cell leukemia. This concept is supported by the observation that donor lymphocyte infusions can be effective in treating B-ALL patients^4^. In addition, several studies have found that one of the strongest predictors of leukemia relapse is the presence of CD4+TIM3+ T cells at diagnosis^5^. These findings, as well as work in murine models of leukemia^6^, suggest that T cell dysfunction plays a role in allowing leukemia relapse. However, treatment of human patients or mice with BCR-ABL+ B-ALL with the checkpoint inhibitor anti-PD1 or anti-PDL1 had no impact on disease progression as a monotherapy^6–8^. In contrast, we recently demonstrated that combination therapy with nilotinib plus anti-PDL1 was very effective at preventing relapse that occurs in ∼90% of mice treated with nilotinib alone^6^. Thus, combination therapy with the TKI nilotinib plus treatment with the checkpoint inhibitor anti-PDL1 (or anti-PD1) is quite effective at preventing relapse in mice treated for B-ALL.

Although nilotinib is approved for treatment of BCR-ABL+ patients, it has been associated with increased arterial occlusive events and is thus used less frequently than a related TKI dasatinib^9^. Dasatinib is associated with other adverse events including platelet dysfunction, but these are viewed as less severe than arterial occlusion^10^. Dasatinib and nilotinib are both second generation, frontline TKIs that circumvent many somatic tyrosine kinase domain (TKD) mutations that confer resistance to imatinib, a first generation TKI^1^. Both dasatinib and nilotinib have greater potency against BCR-ABL^1^ and are associated with better responses than imatinib^11,12^. However, dasatinib has off-target effects on LCK, a SRC-family kinase involved in T cell receptor (TCR) signaling^13^. Thus, a key question is whether dasatinib inhibits anti-leukemia T cell responses. Herein, we compared the effects of dasatinib and nilotinib on T cell function *in vitro* and *in vivo*, and in response to both strong model antigens and potentially less potent leukemia-associated antigens. We found that combination therapy with nilotinib plus anti-PDL1, but not dasatinib plus anti-PDL1, prevented leukemia relapse.

## MATERIALS AND METHODS

### Model

The leukemia cell line LM138, a murine BCR-ABL+ leukemia cell line, was developed by transducing *Arf^−/−^* bone marrow with a retrovirus expressing human BCR-ABL and a GFP reporter^14–16^. *Arf^−/−^* mice were backcrossed to the C57Bl/6 background for >10 generations. All *in vitro* assays were done using T cells from these WT C57BL/6 mice obtained from Charles River Laboratories; *in vivo* studies were also done using T cells from C57Bl/6 mice.

### *In-*vitro assays

#### LM138 TKI IC50 assays

On day 0, GFP+ LM138s were seeded at 2×10^4^ cells in 200 microliters of complete RPMI (RPMI 1640 with 10% fetal bovine serum, 1% penicillin/streptomycin, 1% L-glutamine, 10 mM HEPES, and 50 μM 2-mercaptoethanol). 0.5, 1, 2, 3, 8, 16, or 32 nM of dasatinib or nilotinib, or equivalent volume of solvent control (DMSO) were added to individual wells. Dasatinib (catalog # HY-10181) and nilotinib (catalog # HY-10159) were obtained from MedChem Express (Monmouth Junction, NJ, USA). On day 3, LM138s were counted by flow cytometry (BD LSR II) using antibodies against CD19 (Biolegend catalog #115508, clone 6D5) (San Diego, CA, USA) and counting beads (Invitrogen catalog #PCB100) (Carlsbad, CA, USA), to determine IC50s. We measured percent LM138 survival by dividing the number of LM138s in each well by the average number of LM138s in the control (DMSO) condition. We calculated IC50s through a four-parameter nonlinear regression model, which measures dose inhibition by tuning four parameters (top plateau, bottom plateau, logIC50 and slope) of a curve to the TKI dose vs percent LM138 survival data^17^ (PRISM).

#### T cell proliferation in TKI assays

T cells were isolated from the spleen of a WT mouse and enriched by magnetic depletion using biotin-conjugated depletion antibodies against CD11b (BioLegend, catalog #101204, clone M1/70), CD11c (BioLegend catalog #117304, clone N418), CD19 (BioLegend catalog #115504, clone 6D5), B220 (BioLegend catalog #103204, clone RA3-6B2), GR-1 (BioLegend catalog# 108404, clone RB6-8C5), NK1.1 (BioLegend catalog #108704, clone PK136), F4/80 (Biolegend catalog #123106, clone BM8), and Ter119 (BioLegend catalog #116204, clone TER-119) and Streptavidin microbeads (Miltenyi Biotec catalog #130-048-101) (Bergisch Gladbach, Germany), followed by passage through a Miltenyi LD column (Miltenyi Biotec catalog #130-042-901). Enriched T cells were then labeled with CellTrace Violet (Thermo Scientific catalog #C34557) and were seeded *in-vitro* at 2.5e5 cells/mL in 200 microliters complete RPMI with 100 IU/mL rIL-2 and 0.5 ug/mL anti-CD28 (Tonbo Bioscience catalog #70-0281-U500) and plate-bound anti-CD3 (Tonbo Bioscience catalog #70-0032-U500, clone 17A2) and anti-CD28 (Tonbo Bioscience catalog #70-0281-U500, clone 37.51) (San Diego, CA, USA). After 3 days, T cell proliferation was analyzed by flow cytometry (BD LSRFortessa) using antibodies against CD90.2, NK1.1, Ghost Dye™ Red 780 (antibody details provided in flow panel tables below), and counting beads (Invitrogen catalog #PCB100). Proliferation was measured as the proportion of T cells that had divided more than once, according to CTV intensity.

#### Flow cytometry panels

Leukemia panel (TKI IC-50 assay)

| Target | Fluorophore | Clone | Manufacturer | Catalog number |
| --- | --- | --- | --- | --- |
| CD19 | PE | 6D5 | BioLegend | 115508 |
| GhostDye™Red780 | Red 780 | - | Tonbo Biosciences | 13-0865-T500 |
| BCR-ABL GFP | - | - | - | - |

Leukemia panel (TKI and aPDL1-treated leukemic mice assays)

| Target | Fluorophore | Clone | Manufacturer | Catalog# |
| --- | --- | --- | --- | --- |
| CD19 | BV605 | 6D5 | BioLegend | 115540 |
| GhostDye™Red780 | Red 780 | - | Tonbo Biosciences | 13-0865-T500 |
| BCR-ABL GFP | - | - | - | - |

T cell proliferation panel

| Target | Fluorophore | Clone | Manufacturer | Catalog# |
| --- | --- | --- | --- | --- |
| CD90.2 | FITC | 30-H12 | BioLegend | 105305 |
| NK1.1 | APC-eF780 | PK136 | eBioscience (San Diego, CA, USA) | 47-5941-82 |
| GhostDye™Red780 | Red 780 | - | Tonbo Biosciences | 13-0865-T500 |
| CellTrace™ Violet |  |  | Thermo Scientific | C34557 |

T cell panel (2W/Poly(I:C) mouse immunizations)

Extracellular

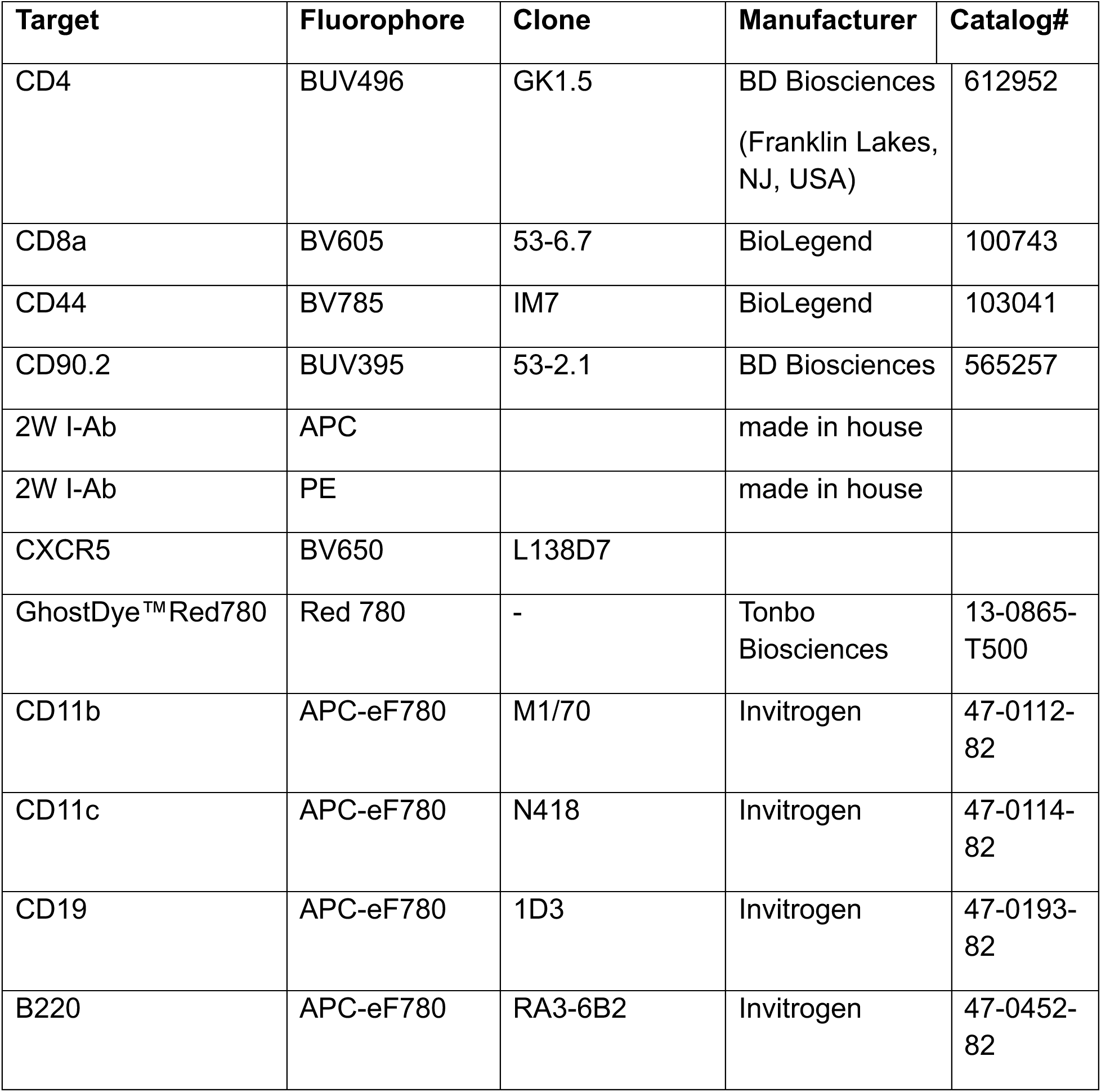
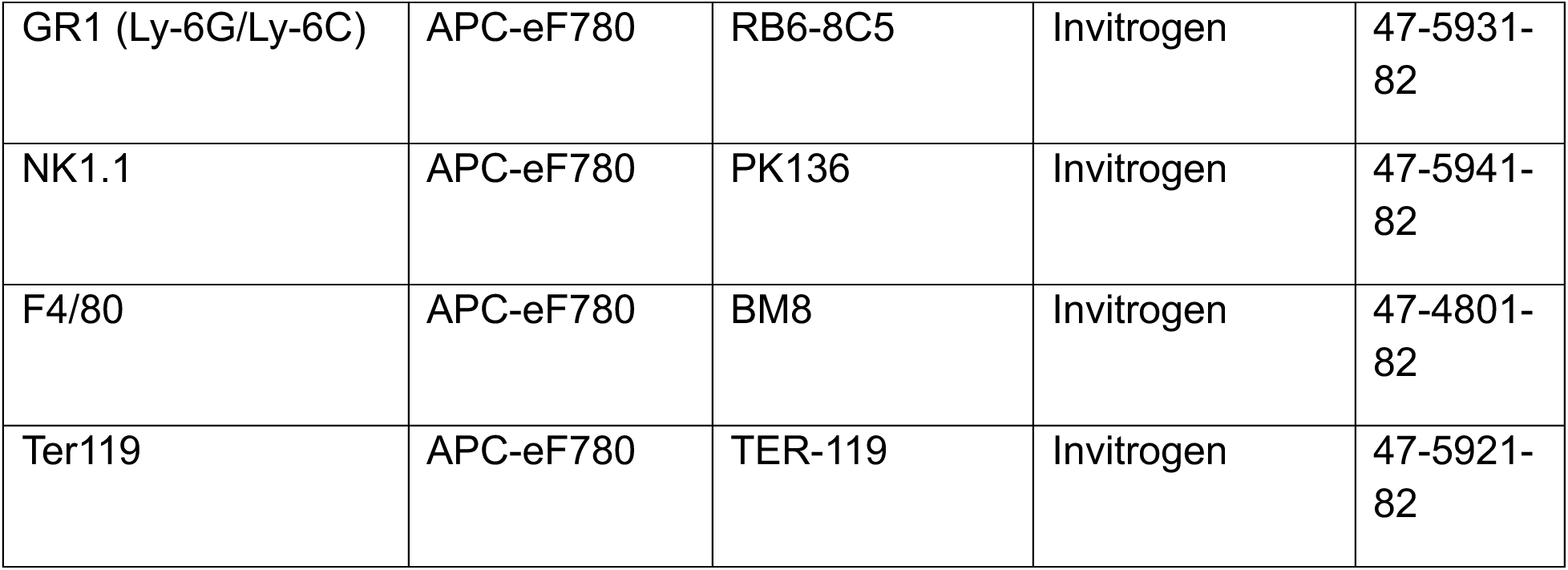

Intracellular

| Target | Fluorophore | Clone | Manufacturer | Catalog# |
| --- | --- | --- | --- | --- |
| Tbet | PE-Cy7 | 4B10 | BioLegend | 644824 |
| FOXP3 | AF488 | FJK-16s | Invitrogen | 53-5773-82 |

T cell panel (TKI/aPDL1)

| Target | Fluorophore | Clone | Manufacturer | Catalog# |
| --- | --- | --- | --- | --- |
| CD90.2 | BUV395 | 53-2.1 | BD Biosciences | 565257 |
| GhostDye™Red780 | Red780 | - | Tonbo Biosciences | 13-0865-T500 |

#### *In-vivo* 2W:Poly(I:C) immunizations

Mice were injected with 100 μL of vaccine containing 20 μg 2W peptide and 100 μg Poly(I:C) in phosphate buffered saline (PBS), intraperitonially as previously described^18^. Mice were treated with 200 μL dasatinib (10 mg/kg/day; Medchem Express catalog #HY-10181), nilotinib (75 mg/kg/dose; Medchem Express catalog #HY-10159), or their respective vehicle controls (citric acid for dasatinib; 10% NMP/90% PEG for nilotinib) by oral gavage on days 0, and 3-6. On day 7, mice were euthanized and their spleens were harvested. To enrich tetramer specific T cells, spleens were incubated with PE and APC labeled 2W:I-Ab tetramers, and later with anti-PE (Miltenyi Biotech catalog #130-048-801) and anti-APC microbeads (Miltenyi Biotech, catalog #130-090-855). Tetramers were designed and produced as previously described^19^. Tetramer-specific T cells were then enriched by passage through a Miltenyi LS column (Miltenyi Biotech, catalog #130-042-401). Samples were analyzed by flow cytometry (BD LSRFortessa) to characterize T cell populations of tetramer positive T cells, using counting beads (Invitrogen catalog #PCB100) to quantify T cell responses.

#### TKI and anti-PD-L1 treated leukemic mouse survival and T cell responses

CD45.2+ C57Bl/6 mice were injected with 2500 LM138s via tail vein. Dasatinib (10 mg/kg/day; Medchem Express catalog #HY-10181), nilotinib (75 mg/kg/day; Medchem Express catalog #HY-10159), ponatinib (5 mg/kg/day; Medchem Express catalog #HY-12047), or solvent control (10% NMP/90% PEG) were administered via oral gavage on days 14-18, and days 21-25. Blocking antibodies against PD-L1 (Bioexcel catalog #BP0101, clone 10F.9G2™) or isotype controls (Bioexcel catalog #BE0088, clone HRPN) (10 mg/kg/dose) were administered on days 14, 16, and 18 intraperitoneally. Mouse survival was recorded to generate survival curves.

#### Early leukemia burden

Early leukemia burden after treatment with TKIs was assessed five days after treatment with TKI or solvent to confirm comparable efficacy in leukemia killing among distinct TKIs. However, some mice were treated with 50 mg/kg/day of dasatinib rather than 10 mg/kg/day. Treatments for all groups were terminated after day 18. On day 19, mice were euthanized, and their spleens were harvested. LM138s were counted by flow cytometry using antibodies against CD19 (BioLegend catalog #115540, clone 6D5) and BCR-ABL and counting beads (Invitrogen catalog #PCB100) to quantify leukemic burdens.

#### T cell expansion

We quantified T cell expansion on day 21. Mice were euthanized and their spleens were harvested. Samples were analyzed by flow cytometry to characterize T cell populations. T cell responses were quantified using counting beads or a hemocytometer.

#### Figures and statistical analysis

Figures were created and statistical analysis was calculated in Graphpad Prism 9. Normality of data was assessed using Shapiro-Wilk tests. Unpaired *t*-tests and Mann-Whitney tests were used to compare means/ranks between two groups when data were normal and not normal, respectively. When data was normal, ANOVA tests were used to compare three or more groups; differences in individual means were assessed using a Dunnett’s multiple comparison test (when comparing to a control group) or a Sidak’s multiple comparison groups (when comparing between different groups). When data was not normal, Kruskal Wallis tests were used instead; differences in medians were assessed using a Dunn’s multiple comparison test. Differences in survival were assessed using Mantel-Cox log-rank test.

## RESULTS

### Dasatinib, but not nilotinib, inhibits T cell proliferation at high doses *in-vitro*

To assess the impact of dasatinib and nilotinib on T cell proliferation, we labeled T cells with Cell Trave Violet (CTV) and activated them with anti-CD3 and anti-CD28 antibodies plus rIL-2 *in vitro* in the presence of varying doses of dasatinib or nilotinib. We measured the percentage of T cells that had divided multiple times by CTV dilution after three days in culture. In separate experiments we treated LM138 leukemia cells with dasatinib or nilotinib and measured the number of LM138s per well on day 3 for each condition relative to controls to determine leukemia cell survival.

The IC50 of dasatinib and nilotinib on leukemia cells was very similar at 5.5 nM and 6.3 nM, respectively (Fig. 1A-B). Dasatinib modestly inhibited T cell proliferation at day 3 *in vitro* at doses >16 nM (Fig. 1C). In contrast, nilotinib had no impact on T cell proliferation at any doses tested (Fig 1D). Thus, dasatinib, but not nilotinib, inhibits T cell proliferation in vitro.

**Figure 1.**
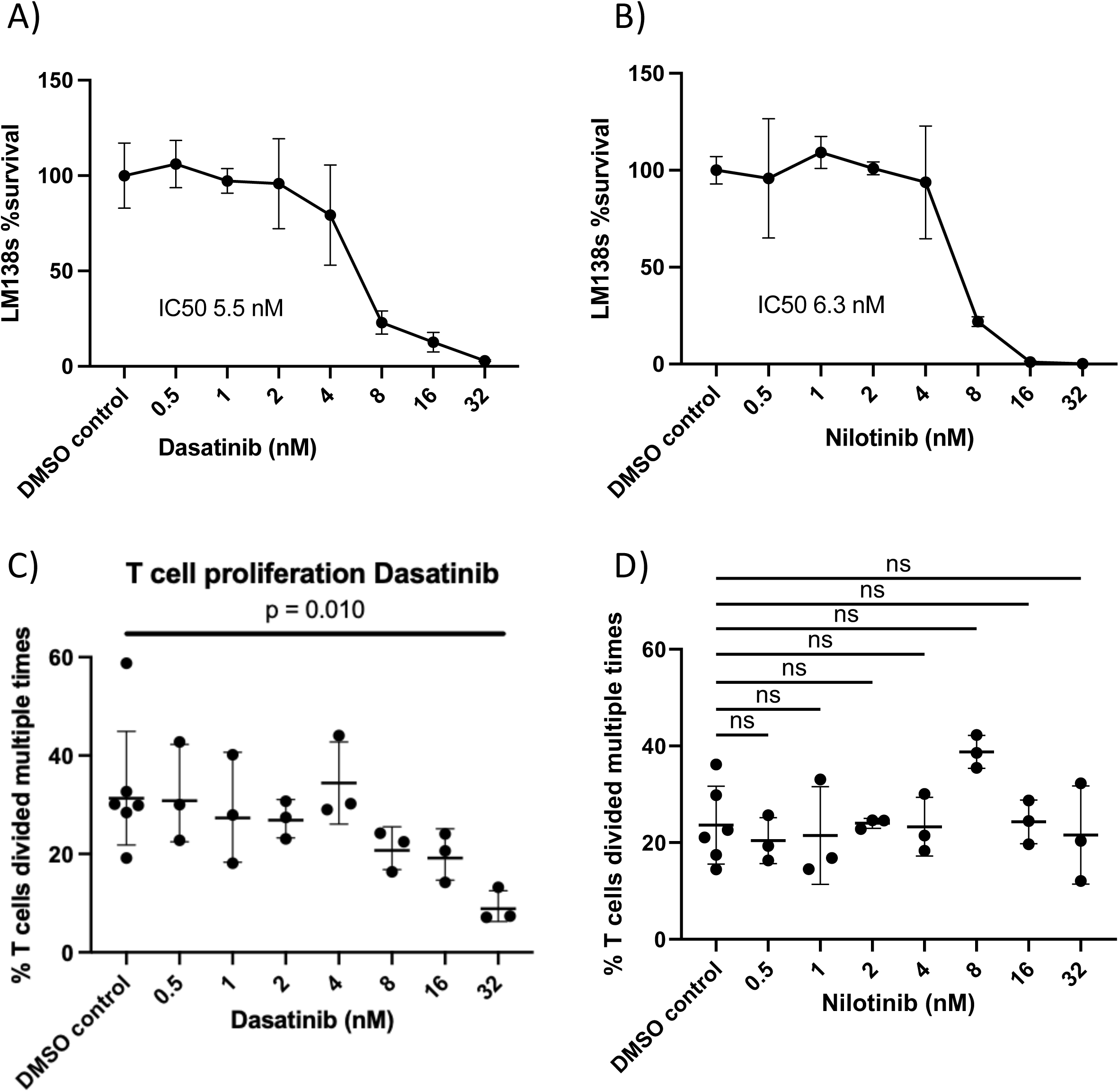
LM138 cells were treated in-vitro with A) dasatinib and B) nilotinib on day 0 and counted on day 3 with counting beads. The proportion of viable cells at different concentrations of each TKI is shown. T cells were isolated from the spleen of a WT mouse, labeled with CellTraceViolet (CTV), and activated in-vitro with aCD3 and aCD28 and cultured with rIL-2. T cells were treated with the indicated concentrations of C) dasatinib and D) nilotinib on day 0 and analyzed on day 3. The proportion of T cells that divided multiple times at different concentrations of each TKI is shown. In C) and D), statistical significance was analyzed using a one-way ANOVA; differences in individual means were assessed using a Dunnett’s multiple comparison test.

### Neither dasatinib nor nilotinib significantly impact CD4 T cell responses to immunization with a strong model antigen in vivo

Because dasatinib inhibited T cell proliferation *in-vitro*, we next investigated whether dasatinib or nilotinib affected T cell responses *in-vivo*. To assess this, we immunized mice with 2W peptide and Poly(I:C) adjuvant, which induces a predominantly CD4 Th1 type T cell response^18^. We treated mice with dasatinib, nilotinib, or their respective solvent controls on day 0, and days 3-6. On day 7, we analyzed the frequency of 2W:I-A^b^ tetramer-specific CD4 T cells. We found that neither the total proliferation of 2W:I-A^b^-specific T cells, nor the relative frequency of CD4+ Th1, Tfh, or Treg T cell subsets were significantly impacted by either TKI relative to controls or untreated mice (Fig. 2A-D).

**Figure 2.**
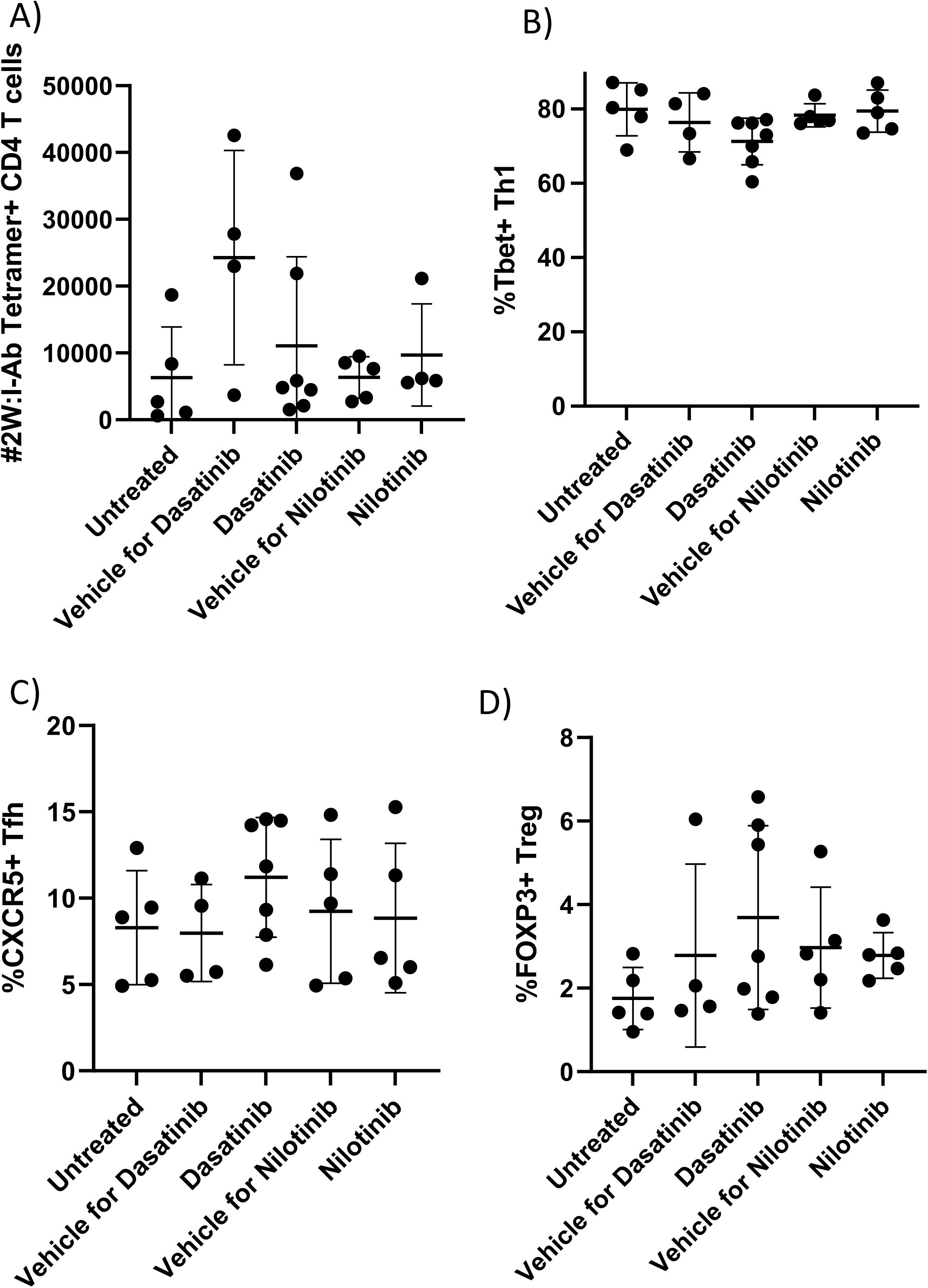
Neither dasatinib nor nilotinib impact CD4 T cell responses to Th type immunization *in-vivo.* WT mice were immunized with 2W/Poly(I:C) intraperitoneally on day 0. Mice were treated via oral gavage with dasatinib, nilotinib, or vehicle controls (citric acid for dasatinib, 10% NMP/90% PEG for nilotinib) on days 0, and 3-6. The mice were sacrificed on day 7, and 2W: I-Ab tetramer positive CD4 T cells were enriched for analysis from the spleen and draining lymph-nodes. A) Total 2W: I-A^b^ Tetramer-positive CD4 T cells in each treatment group. There were no significant differences in means (one-way ANOVA, p = 0.1275). The proportion of B) Tbet+ Th1 cells, C) CXCR5+ Tfh cells, and D) FOXP3+ Tregs among 2W: I-Ab Tetramer-positive CD4 T cells in each group. There were no significant differences among the mean proportions of B) Th1 (p = 0.12 C) Tfh (p = 0.59), or D) Tregs (p = 0.40) as assessed by one-way ANOVA tests, so multiple comparisons were not assessed.

### Nilotinib plus anti-PD-L1 combination therapy confers better survival and increased T cell expansion in leukemic mice than dasatinib plus anti-PD-L1

We previously demonstrated that nilotinib and PD-L1 blockade (which promotes the anti-tumor T cell response) effectively eliminated BCR-ABL leukemia *in-vivo* in >70% of treated mice^6^. Dasatinib is a more broad spectrum TKI and inhibits SRC-family kinases such as LCK, which is required for T cell activation and proliferation^13^. However, dasatinib is also more commonly used to treat BCR-ABL patients. Therefore, we assessed whether the combination of dasatinib plus anti-PD-L1 was as effective as nilotinib plus anti-PD-L1 therapy in preventing leukemia relapse.

For these studies, we injected mice with LM138 leukemia cells, and subsequently treated mice with nilotinib and anti-PD-L1, dasatinib and anti-PD-L1, or vehicle controls and an isotype antibody. We found that leukemic mice treated with nilotinib and anti-PD-L1 had significantly longer survival than mice treated with dasatinib and anti-PD-L1 (Fig. 3A). To confirm that the differences in mouse survival rates were not simply due to differences in the efficiency at which nilotinib and dasatinib repressed leukemia burden at the given doses, we treated leukemic mice with the same doses of TKI, but also treated some mice with 5x the typical dasatinib dose, and measured leukemic burden at day 19 (ie, after the first week of treatment). Both dasatinib and nilotinib were equally effective at eliminating leukemic cells over the first 5 days of treatment (Fig. 3B). Thus, the difference in overall survival observed with combination therapy involving dasatinib versus nilotinib was not due to nilotinib improving clearance of leukemia (if anything dasatinib was slightly better, although differences were not statistically significant). Finally, at day 21, we quantified T cells in the spleen (Fig. 4). We found that mice treated with nilotinib and anti-PD-L1 had significantly greater T cell numbers than mice treated with dasatinib and anti-PD-L1. Thus, dasatinib does have a negative impact on T cell numbers in mice with leukemia.

**Figure 3.**
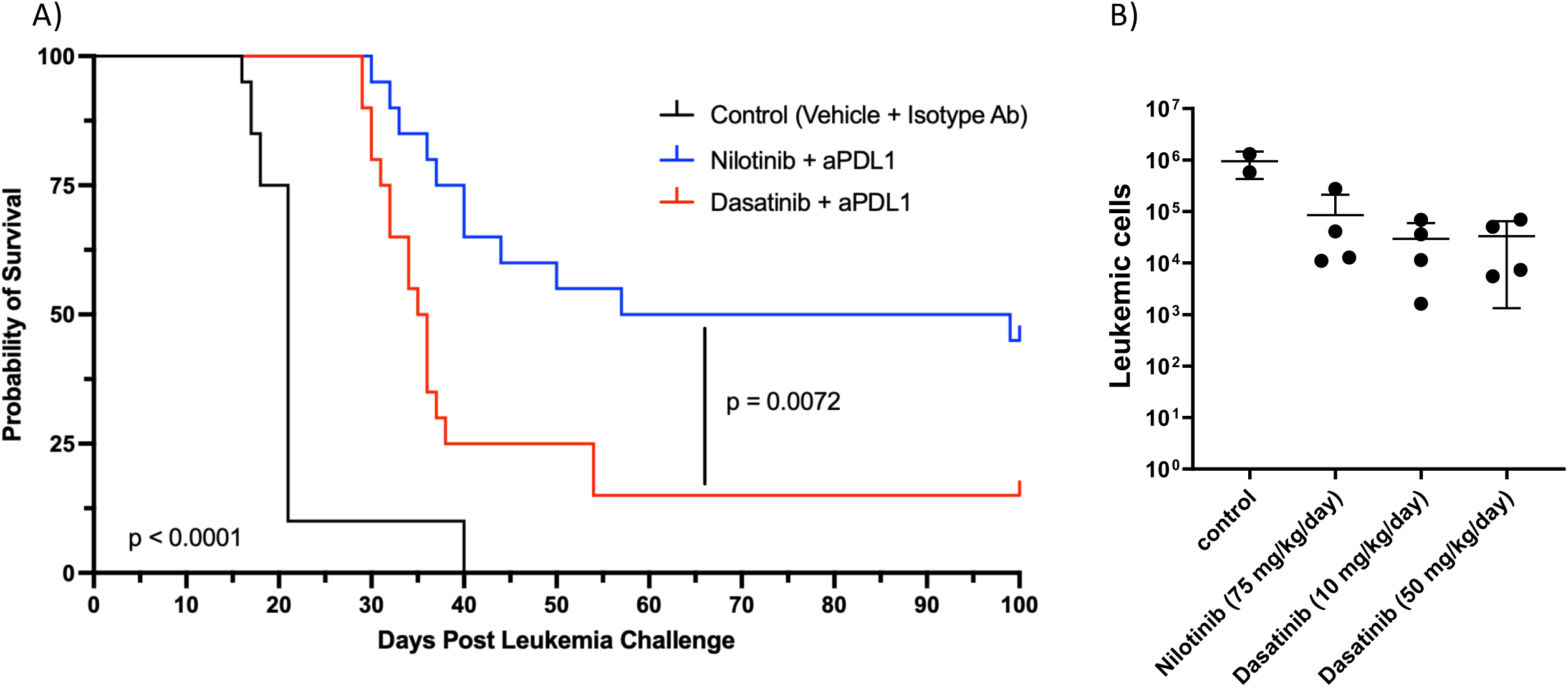
Nilotinib + anti-PD-L1 confers longer survival than dasatinib + aPD-L1 in leukemic mice. CD45.2 mice were injected with 2500 LM138 leukemia cells by tail I.V on Day 0. A) Dasatinib (10 mg/kg), nilotinib (75 mg/kg) or solvent control (10% NMP/90% PEG) were administered by oral gavage days 14-18 and 21-25. Blocking antibodies against PD-L1 or isotype controls (10mg/kg) were administered on days 14, 16 and 18. Survival of the mice in each treatment arm is shown. Statistical significance was analyzed using the Mantel-Cox log-rank test. B) Dasatinib (10 or 50 mg/kg), nilotinib (75 mg/kg), or solvent control (10% NMP/90% PEG) were administered by oral gavage on days 14-18. On day 19, LM138 cells from the spleen were counted. The data is not normal (p = 0.016, Shapiro-Wilk test), so statistical significance was analyzed using the Kruskal-Wallis test. There were no significant differences among the medians (p = 0.16), so multiple comparisons were not assessed.

**Figure 4.**
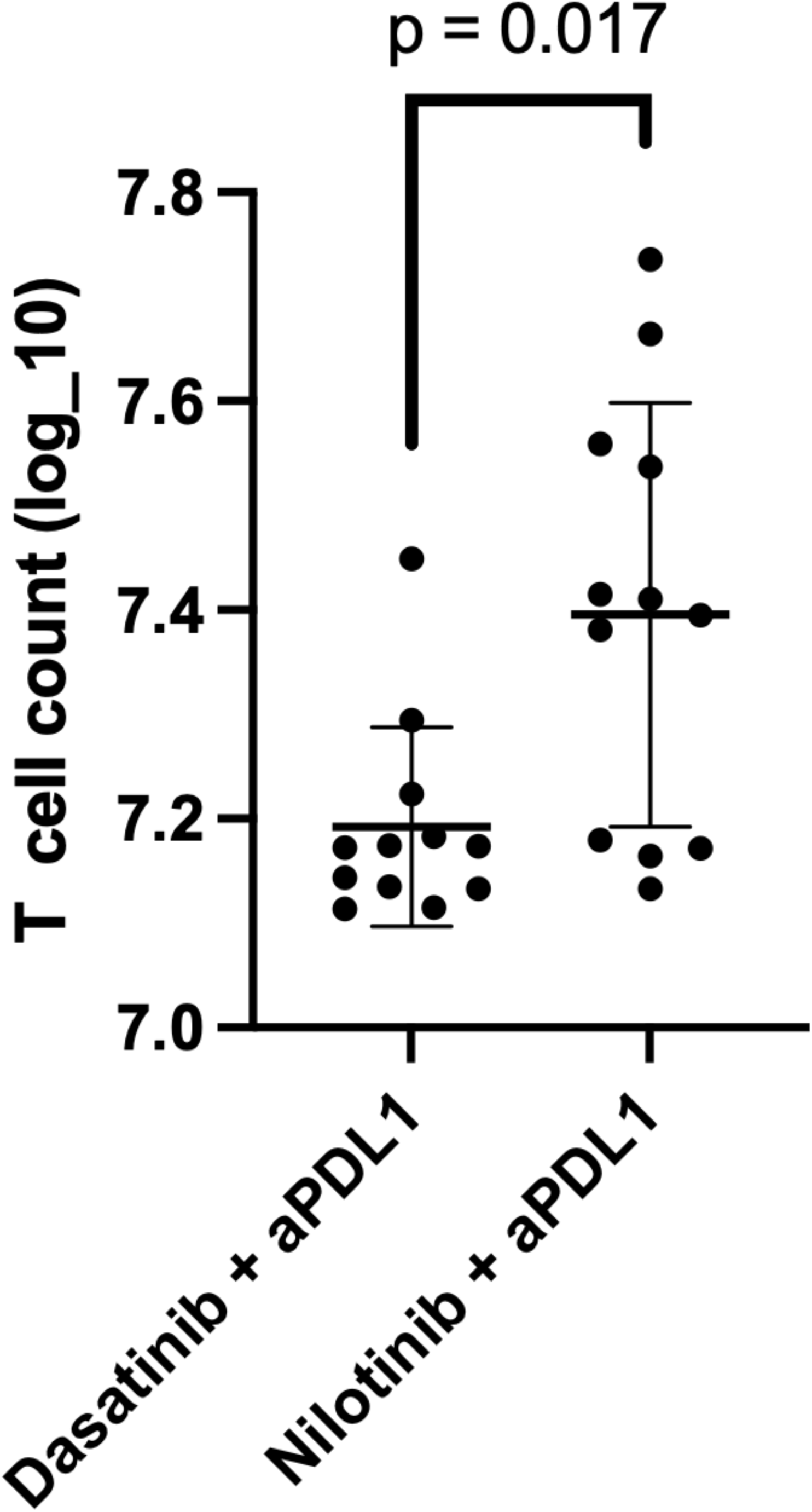
Leukemic mice treated with nilotinib + anti-PD-L1 develop greater T cell expansion than mice treated with dasatinib + aPD-L1. CD45.2 mice were injected with 2500 LM138 leukemia cells by tail I.V on Day 0. Dasatinib (10 mg/kg), nilotinib (75 mg/kg) or solvent control (10% NMP/90% PEG) were administered by oral gavage days 14-18. Blocking antibodies against PD-L1 or isotype controls (10mg/kg) were administered on days 14, 16 and 18. On Day 21, T cells from the spleen were quantified. The data is not normal (Shapiro-Wilk test), so statistical significance was analyzed using the Mann-Whitney test.

To determine whether dasatinib impacted survival even after initial T cell priming we first treated leukemic mice with nilotinib plus anti-PD-L1 for one week followed by dasatinib plus anti-PD-L1 in week two of treatment. To determine whether other SRC family kinase-inhibiting TKIs were also less efficacious than nilotinib in combination with anti-PD-L1, we treated some mice with ponatinib and anti-PD-L1 following injection of LM138 leukemic cells. Ponatinib is less effective at inhibiting SRC family kinases than dasatinib (IC50 of 5 versus 0.5 nM) but both are more potent than nilotinib (IC50>500nM). Mice treated with nilotinib + anti-PD-L1 survived significantly longer than mice treated with sequential nilotinib/dasatinib + anti-PD-1. However, sequential treatment with nilotinib/dasatinib did show a trend towards better survival than treatment with either dasatinib plus anti-PD-L1 or ponatinib plus anti-PD-L1; these latter two treatments were both significantly worse than nilotinib plus anti-PD-L1 (Supplemental figure 1).

## DISCUSSION

The tyrosine kinase inhibitors, dasatinib and nilotinib, are both very effective in treating BCR-ABL leukemia^20^. However, dasatinib, and the third generation TKI ponatinib, also target SRC family kinases^1,13^, while nilotinib does not. As SRC family kinase function is critical for T cell activation, we tested whether dasatinib could inhibit anti-leukemia immune responses. Our findings demonstrate that dasatinib was able to significantly inhibit the ability of the checkpoint inhibitor anti-PD-L1 to prevent relapse in mice treated with dasatinib plus anti-PD-L1, even when nilotinib was used during the first week of treatment, during T cell priming. In contrast, the T cell response to immunization of mice with a potent peptide antigen (2W) in combination with adjuvant was unaffected by dasatinib. The reason for the different response to a leukemia antigen versus a model antigen is unclear. We suggest that one possible explanation has to do with the potency of the peptide antigen and possibly the strength of the adjuvant effect. Leukemia-specific antigens are likely less potent than model antigens and thus T cell responses to those leukemia epitopes may be more susceptible to inhibition by dasatinib. Whether similar effects of dasatinib will be observed for other immunotherapies, such as CAR T cells, is unclear. On the one hand, recent studies suggested that dasatinib helps prevent CD19 CAR T cell exhaustion^13^, and thus would be beneficial for CAR T cell therapy. In contrast, others have reported that dasatinib treatment blocks CD19 CAR T cell activation and function^13^. The reasons for these discrepancies are unclear. In summary, our results indicate that different TKIs can have significant impacts on the subsequent endogenous anti-leukemia T cell response that needs to be considered when designing optimal therapies.

## Acknowledgements

We thank Monica Dinh for assistance with mouse husbandry and Paul Champoux for maintaining the University of Minnesota Flow Cytometry core (supported by P30 CA077598). Enoc Granados Centenos was supported by a Hematology Inclusion fellowship from the American Society of Hematology. This project was supported by an internal grant from the Department of Laboratory Medicine and Pathology and NIH grant R01 AI124512.

## Author Contributions

EM, HV, and EG designed experiments, carried out experiments and analyzed data; GH, TM and LMM assisted with experiments; TD designed and produced MHCII tetramers used in this study; SIT and MAF designed studies and supervised project; EM and MAF wrote the manuscript, all other authors reviewed final manuscript prior to submission.

## Data availability statement

Flow cytometry and Prism files are available upon request.

## REFERENCES

1. Lee, H., Basso, I. N. & Kim, D. D. H. Target spectrum of the BCR-ABL tyrosine kinase inhibitors in chronic myeloid leukemia. Int. J. Hematol. 113, 632–641 (2021).

2. Pfeifer, H. et al. Kinase domain mutations of BCR-ABL frequently precede imatinib-based therapy and give rise to relapse in patients with de novo Philadelphia-positive acute lymphoblastic leukemia (Ph+ ALL). Blood 110, 727–734 (2007).

3. Riva, G. et al. BCR–ABL-specific cytotoxic T cells in the bone marrow of patients with Ph+ acute lymphoblastic leukemia during second-generation tyrosine-kinase inhibitor therapy. Blood Cancer J. 1, e30–e30 (2011).

4. Chauvet, P. et al. Combining blinatumomab and donor lymphocyte infusion in B-ALL patients relapsing after allogeneic hematopoietic cell transplantation: a study of the SFGM-TC. Bone Marrow Transplant. 58, 72–79 (2023).

5. Blaeschke, F. et al. Leukemia-induced dysfunctional TIM-3+CD4+ bone marrow T cells increase risk of relapse in pediatric B-precursor ALL patients. Leukemia 34, 2607–2620 (2020).

6. Tracy, S. I. et al. Combining nilotinib and PD-L1 blockade reverses CD4+ T-cell dysfunction and prevents relapse in acute B-cell leukemia. Blood 140, 335–348 (2022).

7. Cassaday, R. D. et al. Phase 2 study of pembrolizumab for measurable residual disease in adults with acute lymphoblastic leukemia. Blood Adv. 4, 3239–3245 (2020).

8. Manlove, L. S. et al. Heterologous vaccination and checkpoint blockade synergize to induce anti-leukemia immunity. J. Immunol. Baltim. Md 1950 196, 4793–4804 (2016).

9. Aichberger, K. J. et al. Progressive peripheral arterial occlusive disease and other vascular events during nilotinib therapy in CML. Am. J. Hematol. 86, 533–539 (2011).

10. Quintas-Cardama, A., Han, X., Kantarjian, H. & Cortes, J. Dasatinib-Induced Platelet Dysfunction. Blood 110, 2941 (2007).

11. Cortes, J. E. et al. Switching to nilotinib versus imatinib dose escalation in patients with chronic myeloid leukaemia in chronic phase with suboptimal response to imatinib (LASOR): a randomised, open-label trial. Lancet Haematol. 3, e581–e591 (2016).

12. Shen, S. et al. Effect of Dasatinib vs Imatinib in the Treatment of Pediatric Philadelphia Chromosome–Positive Acute Lymphoblastic Leukemia. JAMA Oncol. 6, 358–366 (2020).

13. Weber, E. W. et al. Pharmacologic control of CAR-T cell function using dasatinib. Blood Adv. 3, 711–717 (2019).

14. Manlove, L. S. et al. Adaptive Immunity to Leukemia Is Inhibited by Cross-Reactive Induced Regulatory T Cells. J. Immunol. Baltim. Md 1950 195, 4028–4037 (2015).

15. Manlove, L. S. et al. Heterologous vaccination and checkpoint blockade synergize to induce anti-leukemia immunity. J. Immunol. Baltim. Md 1950 196, 4793–4804 (2016).

16. Williams, R. T., Roussel, M. F. & Sherr, C. J. Arf gene loss enhances oncogenicity and limits imatinib response in mouse models of Bcr-Abl-induced acute lymphoblastic leukemia. Proc. Natl. Acad. Sci. U. S. A. 103, 6688–6693 (2006).

17. Will Prism do a three-parameter or four-parameter logistic fit to my data? GraphPad Software, Inc. https://www.graphpad.com/support/faq/will-prism-do-a-three-parameter-or-four-parameter-logistic-fit-to-my-data/.

18. Kotov, D. I., Kotov, J. A., Goldberg, M. F. & Jenkins, M. K. Many Th Cell Subsets Have Fas Ligand-Dependent Cytotoxic Potential. J. Immunol. Baltim. Md 1950 200, 2004–2012 (2018).

19. Dileepan, T., Malhotra, D., Kotov, D.I., Kolawole, E., Krueger, P.D., Evavold, B.D., Jenkins, M.K. MHC class II teramers engineered for enhanced binding to CD4 improve detection of antgen-specific T cells. Nat Biotechnol 39: 943–048 (2021).

20. Garg, R. J. et al. The use of nilotinib or dasatinib after failure to 2 prior tyrosine kinase inhibitors: long-term follow-up. Blood 114, 4361–4368 (2009).

