## Supplementary Figure 1 for "PDL1 CHECKPOINT BLOCKADE SYNERGIZES WITH NILOTINIB BUT NOT DASATINIB TO PREVENT LEUKEMIA RELAPSE"

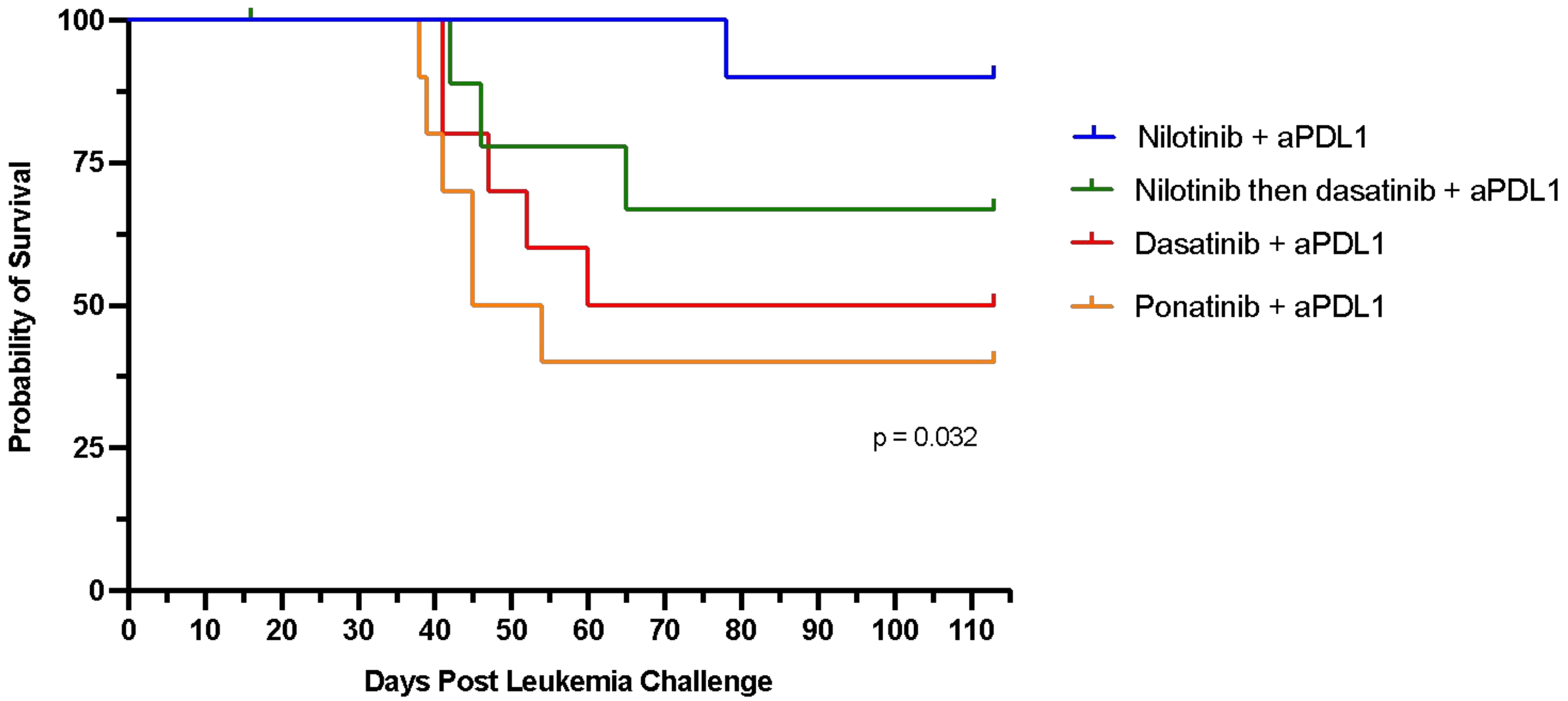


**SUPPLEMENTARY FIGURE 1. Transition from nilotinib to dasatinib later in treatment does not affect the survival of leukemic mice.** Female CD45.2 mice were injected with 2500 LM138 leukemia cells by tail I.V on Day 0. Dasatinib (10 mg/kg), nilotinib (75 mg/kg), ponatinib (5 mg/kg) or solvent control (10% NMP/90% PEG) were administered by oral gavage days 14-18 and 21-25. Alternatively, mice were treated by oral gavage with nilotinib days 14-18 and with dasatinib days 21-25. Blocking antibodies against PD-L1 or isotype controls (10mg/kg) were administered on days 14, 16 and 18. Survival of the mice in each treatment arm is shown. Significance was analyzed using a log-rank test for trend.
